# Anticodon nucleotide modifications affect translational tuning by the ribosomal CAR surface

**DOI:** 10.64898/2026.07.29.740946

**Authors:** Mitsu Raval, Yancheng Zhou, Mia Wichman, Miki Lynch, Daniel Krizanc, Kelly M. Thayer, Michael P. Weir

## Abstract

Nucleotide modifications of the tRNA anticodon can affect protein translation fidelity and speed. Chemical modifications of the anticodon nucleotide 34 are regulated under cellular stress and associated with several translational defects and pathologies. Here, we investigate how these modifications influence A-site codon recognition interactions and their coupling to the CAR site that lies adjacent to nucleotide 34 in the ribosome. The conserved three-residue CAR interface hydrogen bonds in a sequence-dependent manner to the mRNA +1 codon 3’-adjacent to the A-site codon and is implicated in tuning translational speed. The C of CAR is pi-stacked with the nucleotide 34 of the A site tRNA anticodon. The A site and the CAR site influence each other’s hydrogen bonding and stacking interactions, and these codon-adjacency effects potentially provide a layer of regulation affecting translational fidelity and kinetics. Through molecular dynamics simulations of a subsystem of a translocating ribosome IRES-model, we observed that nucleotide 34 modifications affect the hydrogen bonding and stacking interactions at the A site and CAR site as well as CAR’s influence on the A site interactions. Integrating these results with gene sequence and ribosome profiling analyses, we propose that nucleotide 34 modifications help modulate CAR’s sequence-dependent tuning of translation in response to cellular stress.

**Graphical abstract:** 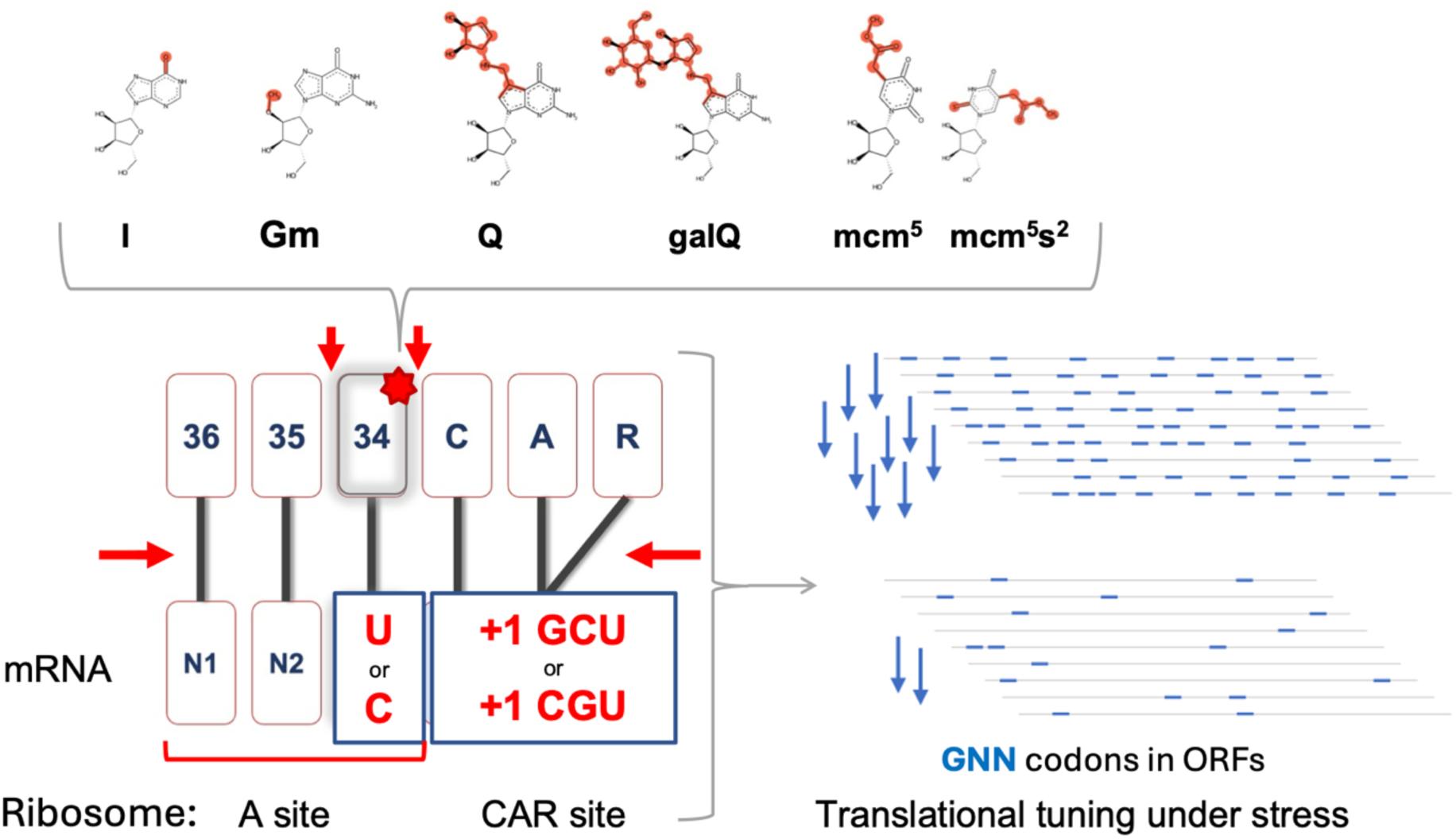

## INTRODUCTION

The CAR interaction surface in the ribosome is a highly conserved motif consisting of C1274 and A1427 of *S. cerevisiae* 18S rRNA (corresponding to C1054 and A1196 in *E. coli* 16S rRNA) and R146 of yeast ribosomal protein Rps3^1^. CAR is located in the small ribosomal subunit facing the mRNA, and immediately adjacent to the ribosome’s A site decoding center (see Fig. 1B). CAR hydrogen bonds (H-bonds) to the mRNA +1 codon 3’ adjacent to the A-site mRNA codon and this interaction is hypothesized to help tune protein translation rates of the ribosome^2,3^. Accurate recognition of the A-site mRNA codon through its H-bonding to the complementary tRNA anticodon nucleotide bases is essential for translation fidelity and efficiency^4–6^. Although protein translation biology has mainly focused on the A-site decoding center in isolation, recent work has suggested the influence of CAR on the A-site base pairing between the codon and anticodon^7^.

**Figure 1:**
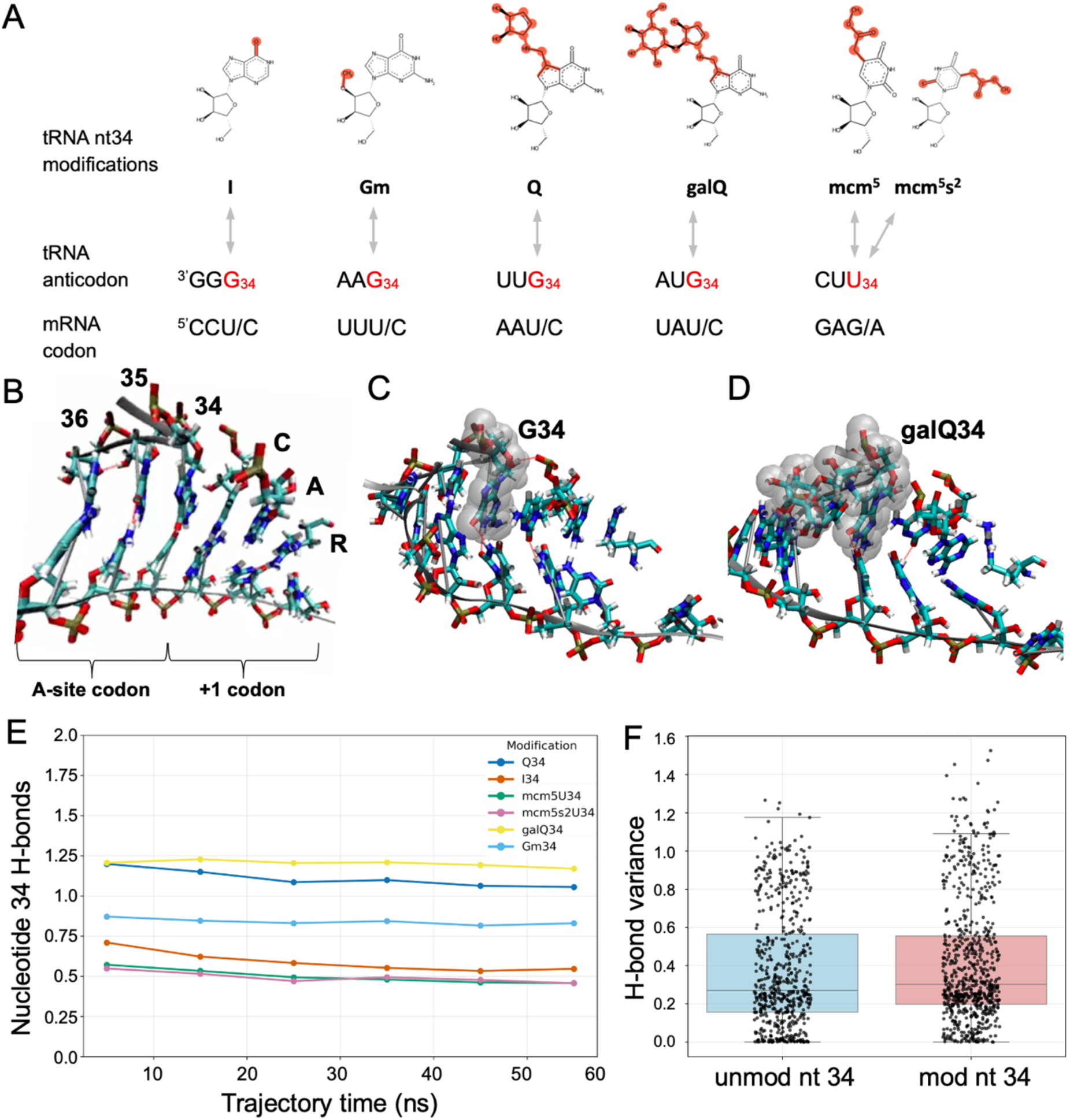
Modification of tRNA nucleotide 34. (A) Nucleotide 34 modifications and corresponding codons and anticodons analyzed in this MD study. (B) CAR is an extension of the A-site anticodon (tRNA nucleotides 34-36). (C, D) Unmodified (C) and galQ modified (D) nucleotide 34 are highlighted with Van der Waals spheres. (E) H-bonding between nucleotide 34 and the wobble nucleotide is shown in a time course of average H-bond counts per frame of MD trajectories with nucleotide 34 modifications. (F) The variances of the H-bond count distributions of MD replicate trajectories for unmodified (unmod) and modified (mod) nucleotide 34 show no significant differences.

The H-bonding of CAR to the mRNA is highly sensitive to the sequence of the +1 codon with which it is interacting. Molecular dynamics (MD) analyses indicate that +1 codons with GNN (where N = A/U/G/C) sequences such as GCU have the strongest H-bond binding to CAR whereas other codons, particularly CGU, demonstrate significantly weaker CAR binding^2^. The MD studies of CAR have made use of a yeast ribosome subsystem at translocation stage II in which a viral internal ribosome entry site (IRES) mimics the mRNA and tRNAs which we refer to for convenience. Of particular interest is A-site tRNA nucleotide 34 which base pairs to the third “wobble” nucleotide of the codon and also pi-stacks with the C nucleotide of CAR, positioning CAR as an extension of the anticodon (see Fig 1). Together, the tRNA anticodon and the pi-stacked residues of CAR align and H-bond with the A-site codon and +1 codon providing an opportunity for dicodon adjacency effects^8^.

Indeed, H-bonding and stacking interactions at both the A and CAR sites are influenced by the sequences of mRNA codons at these sites which together participate in a structural communication network in which CAR is a key regulatory node^7^. Notably, tRNA nucleotide 34 is at the center of this adjacent-site structure. In this study we explore the hypothesis that modified and unmodified chemical structures of nucleotide 34 influence the A and CAR site interactions and the sequence specificity of their translational regulation.

Nucleotide 34 can participate in both wobble base pairing and standard Watson-Crick base pairing, allowing a single tRNA anticodon to recognize multiple mRNA codons^9^. For example, tRNA anticodons with guanosine (G) at nucleotide 34 can bind to A-site codons with uracil (U) or cytosine (C) at the third “wobble” position through wobble or Watson-Crick base-pairing geometries respectively. These differ in their hydrogen bonding capacity as wobble G-U and Watson-Crick G-C base pairs can have up to two and three hydrogen bonds respectively.

Our previous dicodon analysis^3^ of published ribosome profiling experiments with *S. cerevisiae* cells^10^ revealed higher ribosome densities, indicating slower translation speed, for ribosomes mapped to A-site codons with +1 GNN codons. This effect was particularly pronounced for A-site NNU codons that engage in wobble base pairing with tRNA G34. Corresponding MD analyses showed stronger H-bonding between CAR and the +1 codon for +1 GNN codons (+1 GCU compared to +1 CGU). This correlation of stronger CAR-mRNA H-bonding and slower translation supported the model that CAR H-bonding acts as a brake lowering the speed of translation^3^. The CAR brake is regulated by multiple factors. CAR H-bonding is sensitive to the A-site mRNA sequence and nucleotide 34 base pairing geometry, as well as the +1 codon sequence^2,3,7^. Additionally, CAR H-bonding is also sensitive to post-translational modifications like the methylation of arginine (R) or CAR^11^.

Here we explore the influence of chemical modifications of tRNA nucleotide 34 in various mRNA sequence contexts. The nucleotide 34 position is particularly interesting because its base pairing geometry influences CAR H-bonding^3^. Modifications of nucleotide 34 are well studied and biologically relevant (Table 1 and references therein). They are highly regulated under varying environmental conditions like oxidative stress. As these modifications are dynamically regulated under different environmental conditions, nucleotide 34 of the tRNAs effectively store information about the cellular environment in their structures. Moreover, since nucleotide 34 modifications can influence translation fidelity and kinetics, they provide a path for regulating translation in response to environment. The ability of cells to sense and respond to their environment is central to the understanding of cellular evolution and disease manifestation. For example, all of the nucleotide 34 modifications analyzed in this study (Fig. 1A; Table 1) are known to be correlated with neurodegenerative diseases and cancers whose associated clinical pathologies may be understood in part through investigating their tRNA modopathies^12^ and molecular mechanisms impacted by the nucleotide modifications.

**Table 1.**

| **Nt 34**<br>(refs.) * | **Chemical properties** | **Base pairing / structural effects** | **Role in translational regulation** | **Stress regulation** | **Disease Associations** |
| --- | --- | --- | --- | --- | --- |
| I (9,13–25) | A→I<br>(deamination;<br>C6 NH<sub>2</sub>→O) | Expands base pairing to U, C, A; flexible wobble geometry | Expands decoding capacity; modulates elongation kinetics | ADAT2/3-mediated; essential for viability | Neurodevelopmental disorders, myositis, intellectual disability |
| Gm (26–44) | Ribose 2'-OH→OCH<sub>3</sub> | Increased rigidity; stabilizes sugar pucker | Enhances decoding stability; prevents ribosome pausing | Upregulated under oxidative stress (Trm7, FTSJ1 mediated) | Intellectual disability, cancer metabolism, neuronal defects |
| Q (45–65) | Bulky cyclopentenediol on 7-deazaguanine | Reduced flexibility; reduces differential | Reduces codon bias; improves fidelity; stabilizes tRNA | Nutrient-dependent (microbiome-derived queuine) | Cancer, neurodevelopment, gut disorders |

|  |  | NNC/NNU translation |  |  |  |
| --- | --- | --- | --- | --- | --- |
| galQ<br>(45,47,55,66) | Q + galactose sugar | Increased steric bulk and polarity | Tunes codon bias; suppresses stop codon readthrough | Also depends on queuine availability | Similar to Q; emerging physiological roles |
| mcm<sup>5</sup>U<br>(67–70) | 5-methoxy-carbonylmethyl addition | Electron-withdrawing; increases flexibility | Enables wobble decoding (NNG codons); supports elongation | Regulated by Elongator/Trm9; stress-responsive | Cancer, stress response defects |
| mcm<sup>5</sup>s<sup>2</sup>U<br>(38,67–75) | mcm5 + 2-thio substitution | Increased rigidity; s<sup>2</sup> favors NNA codon base pairing | Maintains reading frame; improves fidelity; prevents stalling | URM1 pathway; regulated by ROS and nutrients | Neurodegeneration, mitochondrial disease, DREAM-PL syndrome |

\* Nucleotide 34 modifications studied here are Inosine (I34), 2'-O-methyl guanosine (Gm34), queuosine (Q34), galactosyl queuosine (galQ34), 5-methoxycarbonylmethyluridine (mcm<sup>5</sup>U34) and 5-methoxycarbonylmethyl-2-thiouridine (mcm<sup>5</sup>s<sup>2</sup>U34).

In this study, we used molecular dynamics (MD) simulations of the ribosome’s decoding center neighborhood to explore the effects of nucleotide 34 modifications on H-bonding and pi-stacking at the A site and CAR site. In addition to direct effects on nucleotide 34 base pairing and pi-stacking, we also observed effects across the full A-site codon/anticodon as well as effects on CAR stacking and H-bonding with the +1 codon. Of particular note were observations that one of the tested modifications (2′-O-methyl guanosine) increases the dynamic range of CAR H-bonding sensitivity to the +1 codon sequence suggesting that nucleotide 34 modifications can influence CAR tuning of translation.

## METHODS

To obtain a comprehensive view of the sequence-specific effects of the environment-sensitive modifications of tRNA nucleotide 34 on the CAR-mediated translational regulation, we integrated insights from molecular dynamics simulations and bioinformatics analyses. We analyzed various constructs of a ribosome decoding center neighborhood with different nucleotide 34 modifications, base pairing geometries and CAR site +1 codons. Relationships between CAR site sequence specificity and dicodon genomic frequencies and translation rates were assessed to investigate the potential roles of CAR in tuning protein translation.

### Molecular Dynamics Simulations

Molecular dynamics (MD) simulations of biomolecular structures predict atomic motions based on force fields that rely on the laws of physics^76^. Our all-atom MD simulation structure coordinates were derived from the cryo-EM structure of an *S. cerevisiae* ribosome at translocation stage II (PDB ID 5JUP)^77^, a stage when CAR exhibits strong H-bonding to the +1 codon. As described previously, we simulated a 494 residue subsystem centered around the tRNA nucleotide 34 and the adjacent C of CAR with the outer residues (onion shell) restrained to maintain the translocation stage II conformational state^11^. Nucleotide substitutions were introduced in this starting structure using Amber22 tLEAP as described previously^2^.

To add RNA modifications, we edited the unmodified nucleotide 34 residue rows in the PDB structure files. We retained the coordinates of all the atoms common in the unmodified and modified residues, deleted rows of other atoms, edited the residue identity (e.g. G to Gm) and grew the remaining atoms in tLEAP. Proteins, RNA, water and modifications were parameterized using ff14SB, OL3, TIP3P, modrna08, prepin, and frcmod force fields respectively. Next, structures were heated, equilibrated, simulated and analyzed using our established protocols that have been described previously^2,11^. MD replicates of each structure were generated by independently assigning random velocities to atoms from a gaussian distribution corresponding to a temperature of 300K.

Similar to molecular genetics experiments, we performed computational genetics experiments in which we introduced nucleotide modifications and substitutions and examined how they altered the resulting H-bonding and stacking phenotypes. Different combinations of the nucleotide identities and post-transcriptional modifications were tested with particular attention paid to three variables: (i) the modification of nucleotide 34 and its associated codon/anticodon, (ii) the base pairing geometry (wobble or Watson-Crick) of nucleotide 34, and (iii) the +1 codon sequence (GCU or CGU) that H-bonds with CAR.

For the first variable, we analyzed six different tRNA nucleotide 34 modifications outlined in Table 1: (i) inosine (I34) formed by adenine deamination which allows base pairing with A, U or C; (ii) 2′-O-methyl guanosine (Gm34), a methylation of the G34 ribose which increases structural rigidity; (iii) queuosine (Q34), a bulky cyclopentenediol ring that increases steric hinderance; (iv) galactosyl queuosine (galQ34) where the galactose sugar addition to Q further increases the steric bulk and adds more polarity; (v) 5-methoxycarbonylmethyluridine (mcm^5^U34) whose mcm^5^ group is electron withdrawing and adds flexibility; and (vi) 5-methoxycarbonylmethyl-2-thiouridine (mcm^5^s^2^U34) where the thiolation adds polarity. *In vivo*, these modifications are found on specific tRNAs with a particular tRNA anticodon and corresponding amino acid^78^. We substituted A-site tRNA anticodon and mRNA codon nucleotides so that these modifications were analyzed with one of their native A-site anticodon sequences (Fig. 1A, Table S1).

Modified nucleotide 34 structures were compared with corresponding structures with unmodified residues. However, I34 (derived from adenosine) is only present in eukaryotes and was compared to its G34 prokaryotic counterpart^14,15^. Gm34, Q34 and galQ34 are all derivatives of G34 which all can bind to both U and C of A-site NNU and NNC codons in wobble and Watson-Crick base pairing geometries respectively (the second tested variable)^9^. In contrast, U34, mcm⁵U34 and mcm⁵s^2^U34 anticodons can bind to mRNA codons NNG and NNA with wobble and Watson-Crick base pairing geometries respectively and were structured correspondingly in our simulations. For the third tested variable, the +1 codon sequence, we used +1 GCU and +1 CGU which exhibit strong and weak H-bonding to CAR respectively^2^.

We used different combinations of the three variables in our MD analysis giving 44 starting structures (summarized in Table S1). For all 44 structures, each with a unique combination of A-site, +1 codon and nucleotide 34 modification, we generated 30 independent MD replicates (20 × 60ns and 10 × 100ns long trajectories). RMSD values of the trajectories converged to a plateau within the first 20ns of dynamics (Fig. S1). Hence, we restricted out analyses to after 20ns of dynamics. For each trajectory, we measured average H-bonding across all simulation frames by characterizing binary values for H-bonding occurrences using the default distance (3.5 Å) and angle (135°) cutoffs between the atoms of participating residues. This was done using the Hbond function in cpptraj^79^. Statistical analyses were performed using Levene’s test and student’s t-tests for comparison of variances using data from all frames or 30 replicate averages respectively^80^.

### Dicodon Analysis

Dicodon frequencies were measured in high-expression yeast ribosomal protein genes to investigate possible sequence biases consistent with CAR-mediated tuning of translation. Sequence walkers were used to categorize UUU (TTT) codons with or without 3’-adjacent GNN codons. The fractions of UUUGNN dicodons in ribosomal protein gene ORFs in multiple species were compared with corresponding fractions for all genes. Bootstrap analysis was performed selecting from all genes with replacement using the number of ribosomal protein genes as the sample size. For each species, the observed UUUGNN dicodon ratio was compared with the bootstrap distribution (10,000 samples) to assess statistical significance.

### Ribosome Profiling Analysis

Published ribosome profiling data^10^ were analyzed for dicodon frequences as described previously^3^. Ribosome profiles for *S. cerevisiae* cells with and without oxidative stress treatment (0.03% H₂O₂) were compared (two experiments for each growth condition). Ribosome footprints were mapped to mRNA sequences using A-site offsets based on footprint length and the peaks from the periodic three-nucleotide patterns were used for analysis.

Ribosome footprint densities for each codon type were normalized to the other codons in the same gene’s ORF ensuring that each gene received equal weighting in the analysis. Genes with ORFs >200 nucleotides and >0.1 footprint per nucleotide were included in the analysis which was restricted to footprints in the first 200 nucleotides of ORFs. Bootstrap analysis was performed as described previously^3^ by sampling genes with replacement from non-stress experiments and using the resulting bootstrap distributions to assess dicodon ribosome densities from stressed cells.

## RESULTS AND DISCUSSION

To test the hypothesis that modifications of tRNA nucleotide 34 influence A-site codon recognition and CAR braking interactions, we examined the effects of a group of six nucleotide 34 modifications on MD behavior of a subsystem of the A-site decoding center (Fig. 1A, Table 1). The modifications that were examined were inosine (I34), 2′-O-methyl guanosine (Gm34), queuosine (Q34), galactosyl queuosine (galQ34), 5-methoxycarbonylmethyluridine (mcm⁵U34), and 5-methoxycarbonylmethyl-2-thiouridine (mcm⁵s²U34). We edited the MD subsystem to examine modifications with their associated A-site codons and anticodons (as described in Methods) in the context of 3’-adjacent (+1) codons +1 GCU and +1 CGU which exhibit strong and weak CAR braking behaviors respectively (Fig. 1, Table S1). For ease of presentation, we henceforth refer to these modifications with the first two nucleotides of their corresponding A-site codon. For example, nucleotide 34 inosine was simulated with A-site codons CCU and CCC and is referred to as I34_CC.

### Nucleotide 34 modifications affect A-site base pairing

Since modifications introduced chemical perturbations in the system, to ensure that the modifications were incorporated stably, and our structures were equilibrated globally, we analyzed backbone atom RMSD values of all structures which had settled into a plateau after 10-20 ns (Fig. S1). Since some of the nucleotide 34 modifications were quite bulky (Fig. 1C, D), we ensured that their incorporation was stable and that they did not induce abnormal structural behaviors by testing the MD behaviors over time. We examined time courses of H-bonding of nucleotide 34 to its base-pairing partner, the third wobble nucleotide of the A-site codon. This revealed stable H-bonding over time reflecting post equilibration dynamics (Fig. 1E).

Additionally, we tested if the modifications altered the fluctuation of H-bonding over time relative to their unmodified counterparts by examining the variance of nucleotide 34 H-bond count data distributions of each MD replicate across 0-6000 frames (corresponding to 0-60 ns of trajectory time) (Fig. 1F). Levene’s test (p = 0.995) showed that the time course variance of the replicate frames of unmodified (nucleotide 34) trajectories was not significantly different compared to the modified trajectories. Thus, the modifications were successfully incorporated and did not destabilize the structures. These results were consistent with backbone atom RMSD time courses which showed settling (convergence) after 10-20 ns (Fig. S1). Correspondingly, our analysis described below is based on MD trajectories of 20-60 ns (20 MD replicates) and 20-100 ns (10 replicates).

To assess A-site codon : anticodon interactions, we first quantified the H-bonding levels between tRNA nucleotide 34 and the third “wobble” nucleotide of the A-site codon. This base pairing can exhibit wobble or Watson-Crick geometries. For G34, G:U (wobble) and G:C (Watson-Crick) base pairs can form up to 2 and 3 H-bonds respectively, whereas for U34, U:A and U:G both form up to 2 H-bonds. For G34, our analysis consistently showed higher H-bonding levels for Watson-Crick base pairing as expected (Fig. 2). This consistency of our results with the known nucleotide chemistry behaviors helped validate our simulation and analysis methodology. As confirmed by visualization of trajectory frames, the observed H-bonding was highly dynamic showing each H-bond in some but not all frames thus accounting for the average H-bonding measurements being lower than the theoretical maxima. We also note that the variation in positioning of the wobble nucleotide also led to occasional H-bonding with tRNA (anticodon) nucleotide 35 or C of CAR (which were not included in the H-bonding measurements in Figure 2).

**Figure 2:**
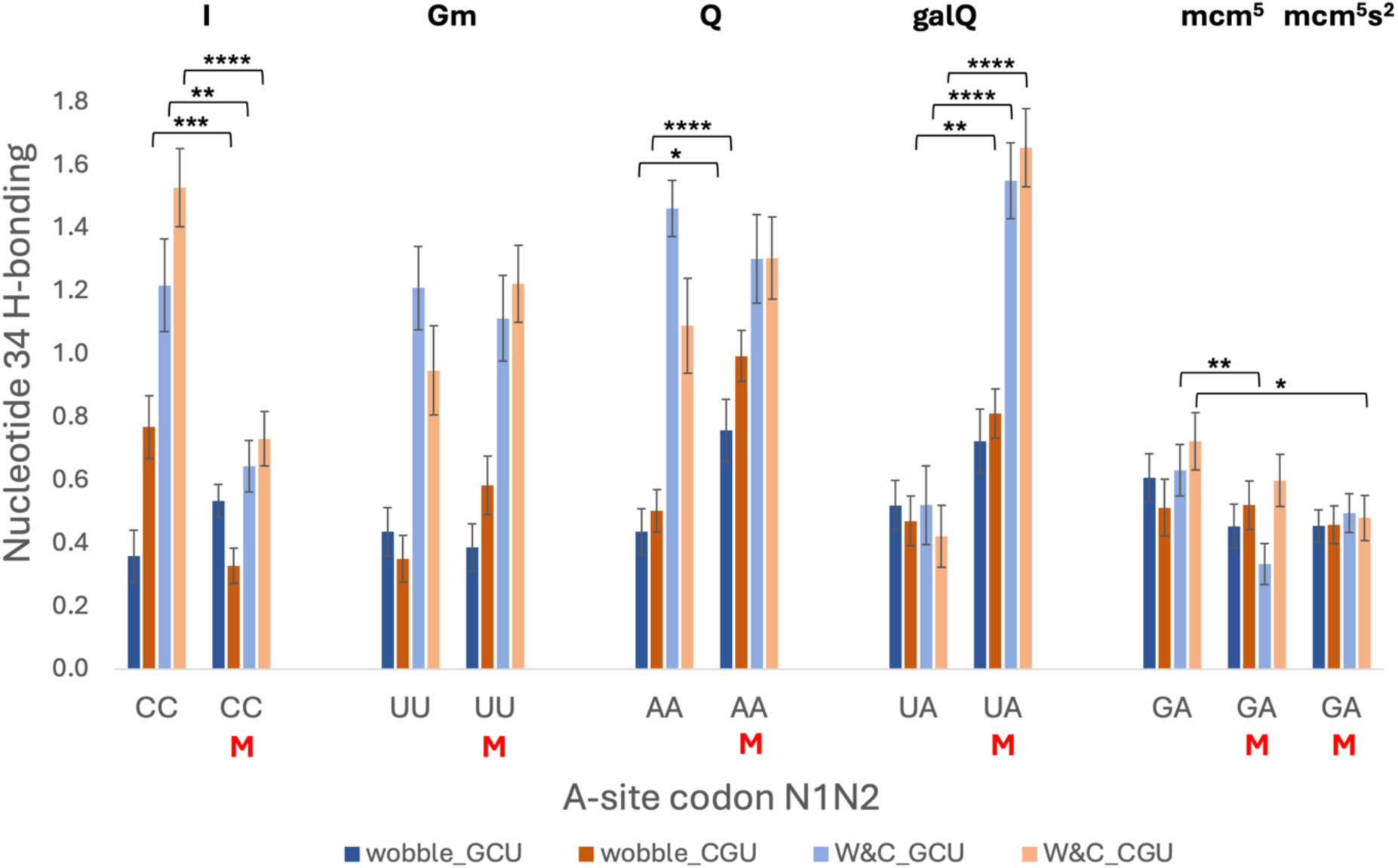
Modifications alter nucleotide 34 base pairing. H-bonding between nucleotide 34 and its base-pairing partner A-site codon nucleotide 3 was averaged across all the frames of 30 MD replicates for each structure. The x-axis (this figure and Figs. 3, 4 and 5) shows the first two nucleotides (N1N2) of the A-site codon which were tested with wobble and Watson-Crick base pairing of the third nucleotide (see Fig. 1A) and with +1 codons GCU or CGU. Bars corresponding to the modified structures are indicated (M; also for Figs. 3, 4 and 5). Student’s t-test comparisons between unmodified and modified nucleotide 34 H-bonding. p-values are denoted as: * p<0.05, ** p<0.01, *** p<0.001, **** p<0.0001, ns p≥0.05 (no * sign).

We next assessed the effects of modifications on nucleotide 34 base pairing to the A-site nucleotide three (wobble position). Comparisons of this base pairing with and without the nucleotide 34 modifications showed differences in H-bonding which were significant for the I34_CC, Q34_AA, galQ34_UA and mcm^5^U34_GA (Fig. 2). Thus, these chemical perturbations of nucleotide 34 can alter its base pairing properties.

Given the effects of nucleotide 34 modifications on base pairing at the third codon position (Fig. 2), we expanded our analysis to examine effects over the full A-site codon (Fig. 3). Accordingly, we measured the sum of H-bonding between all three A-site mRNA codon nucleotides with the three complementary tRNA anticodon nucleotides. This broader measurement of H-bonding included the rarer cases of nucleotide base pairing with their off-frame partners (e.g. wobble nucleotide with tRNA nucleotide 35) as well as cases of base pairing with C of CAR. The resulting assessment revealed significant decreases and increases in A-site H-bonding for I34_CC and galQ34_UA respectively relative to their unmodified G34 controls.

**Figure 3:**
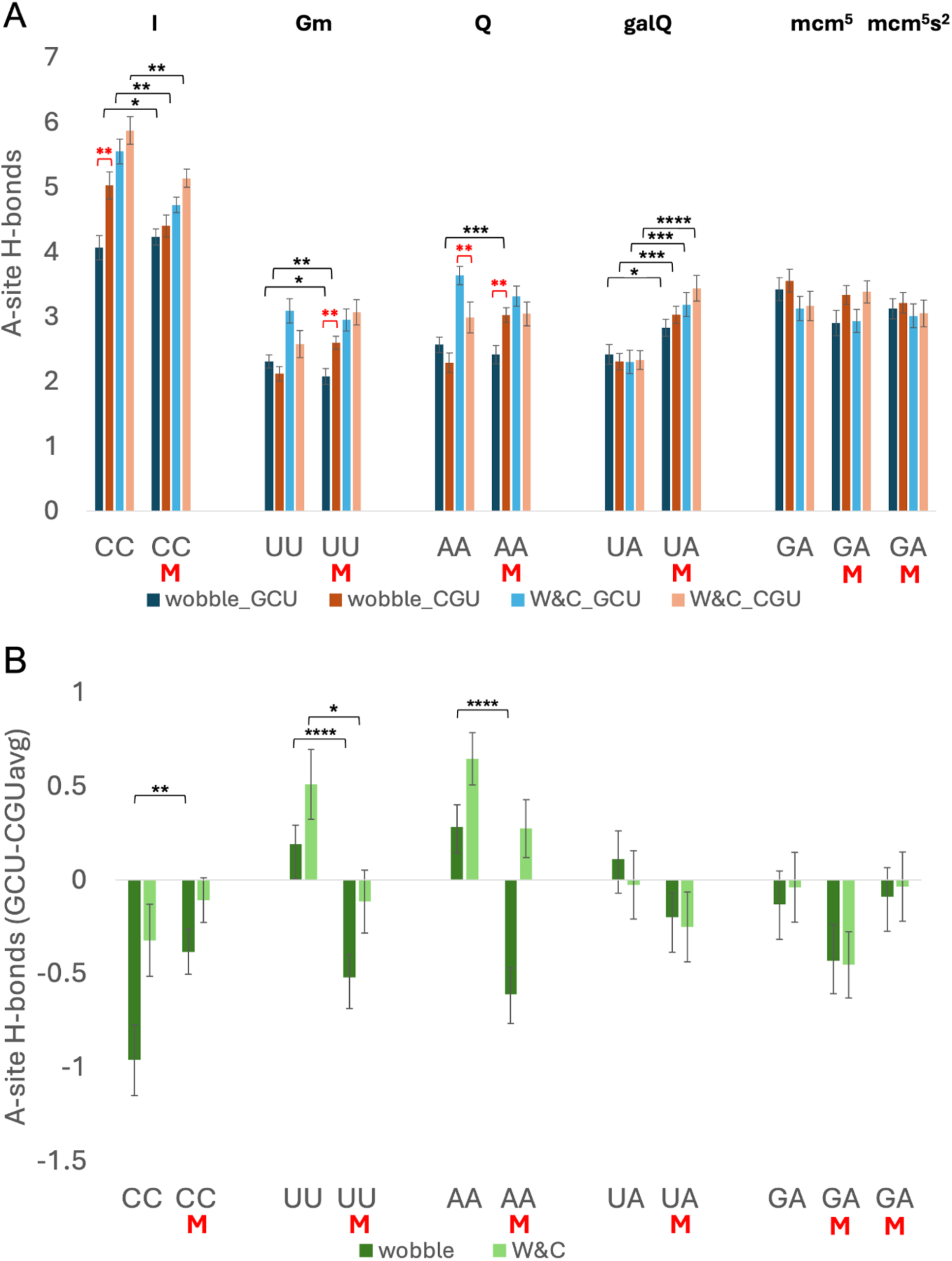
Modifications affect A-site H-bonding. (A) H-bonding by the A-site codon was calculated by summing each mRNA nucleotide’s H-bonding with its in-frame and (occasional) -1 and +1 frame partners (e.g. wobble nt H-bonding with tRNA nt35+nt34+C of CAR). Average values across 30 MD replicates were plotted for each structure. (B) The A site’s CAR sensitivity calculated by subtracting the average A-site H-bonding of all 30 +1 CGU replicate simulations from the A-site H-bond count for each individual +1 GCU MD replicate giving a distribution of 30 GCU-CGUavg difference values. The means and standard errors of these distributions are plotted. Student’s t-test p-values are denoted as: * p<0.05, ** p<0.01, *** p<0.001, **** p<0.0001, ns p≥0.05 (no * sign).

While I34_CC and galQ34_UA modifications showed a clear statistically significant change for all A-site codon contexts, Gm34_UU and Q34_AA showed interesting context specific effects (Fig. 3A, black brackets). Their A-site H-bonding increased or decreased depending on whether the CAR site +1 codon sequence was GCU or CGU. For example, for wobble (dark blue and red) UUU codon context, Gm34_UU decreases the A-site H-bonding when the +1 codon sequence is GCU (dark blue) but increases it in the presence of adjacent +1 codon sequence CGU (dark red). Similarly, for wobble (dark blue and red) AAU codon context, Q34_AA does not change the A-site H-bonding with +1 GCU (dark blue) but significantly increases it in the presence of +1 CGU (dark red).

To emphasize the effects of the CAR site on the A-site codon : anticodon H-bonding, statistical comparisons of these H-bonding values between +1 GCU (blue shades) and +1 CGU (red shades) structures were made using student’s t-tests (Fig. 3A, red brackets). H-bonding is significantly different for +1 GCU and +1 CGU for many cases i.e., G34_CCU, Gm34_UUU, G34_AAC, Q34_AAU. Thus, as previously reported, A-site H-bonding is sensitive to the CAR site sequence^7^. Of particular interest in this study, two of these differences were only observed for modified versions of nucleotide 34. For the Gm34_UUU and Q34_AAU structures, A-site H-bonding was significantly higher for CAR site +1 CGU compared to +1 GCU, but this difference was not observed without the nucleotide 34 modifications (Fig. 3A).

To systematically interpret how modifications alter CAR’s influence on the A site, we defined a metric to quantify CAR sensitivity of the A site, the magnitude of change in A-site H-bonding that we observed after altering the +1 codon sequence at the CAR site. To quantify A site’s CAR sensitivity, we compared A-site H-bonding in +1 GCU and +1 CGU contexts by measuring A-site H-bonding GCU-CGUavg. To calculate GCU-CGUavg we took the 30 values for A-site H-bonding measured in the 30 +1 GCU MD replicates and subtracted from each the average A-site H-bonding of the 30 +1 CGU replicates 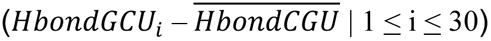 We observed that the A site’s CAR sensitivity is modulated by the modifications I34_CCU, Gm34_UUU, Gm34_UUC and Q34_AAU (Fig. 3B). Nucleotide 34 modifications can either increase (UUU, AAU) or decrease (CCU, UUC) the magnitude of the sensitivity, and in these cases, the +1 GCU with modification can either lead to higher or lower H-bonding at the A site depending on the A site context.

In summary, nucleotide 34 modifications can alter A-site codon : anticodon H-bonding. The magnitude and the directionality (increase or decrease in H-bonding) of these effects can be tuned by the adjacent CAR site +1 codon sequence given that modifications impact the A site differently for +1 GCU and +1 CGU contexts (Fig. 3A; black brackets). Similarly, the effects of the CAR site on the A site are also tuned by nucleotide 34 modifications such that the difference between H-bonding with +1 GCU and +1 CGU are significantly altered by the modifications (Fig. 3A; red brackets). Together these results support a model of structural interplay between the A site, CAR site and nucleotide 34 modifications where the CAR site tunes the effect of the modifications on the A site (Fig. 3A; black brackets) and the modifications tune the effects of the CAR site on the A site (Fig. 3A; red brackets). This underscores a model of translational regulation where A-site codon recognition functionality and therefore, translational fidelity are influenced by the structural H-bonding effects of nucleotide 34 modifications, the CAR site, and their interplay.

### Pi-pi stacking is modulated by nucleotide 34 modifications

Following the analysis of how modifications affect H-bonding of the A-site codon and anticodon, we also examined their effects on base stacking of nucleotide 34 (Fig. 4). Pi-pi stacking of nucleotide 34 with nucleotide 35, its 3’ anticodon partner, was measured by monitoring changes in the distance between the centers of geometry (COG) of the two nucleotide base rings participating in the stacking^2,3^ (Fig. 4A). Lower COG values reflect better stacking, and based on systematic analysis of stacked structures^81^, COG distances less than 4.5Å were considered to reflect well stacked bases. While many of the stacking interactions of nucleotide 34 were not significantly affected by modifications of the nucleotide, we observed several cases of significant effects of modifying nucleotide 34. When we assessed the 35 : 34 stack, within the tRNA anticodon, we found that the addition of galactosyl queuosine modification onto nucleotide 34 (galQ34_UA) significantly improved its stacking with nucleotide 35 in the four tested mRNA sequence contexts (Fig. 4A, black brackets).

**Figure 4:**
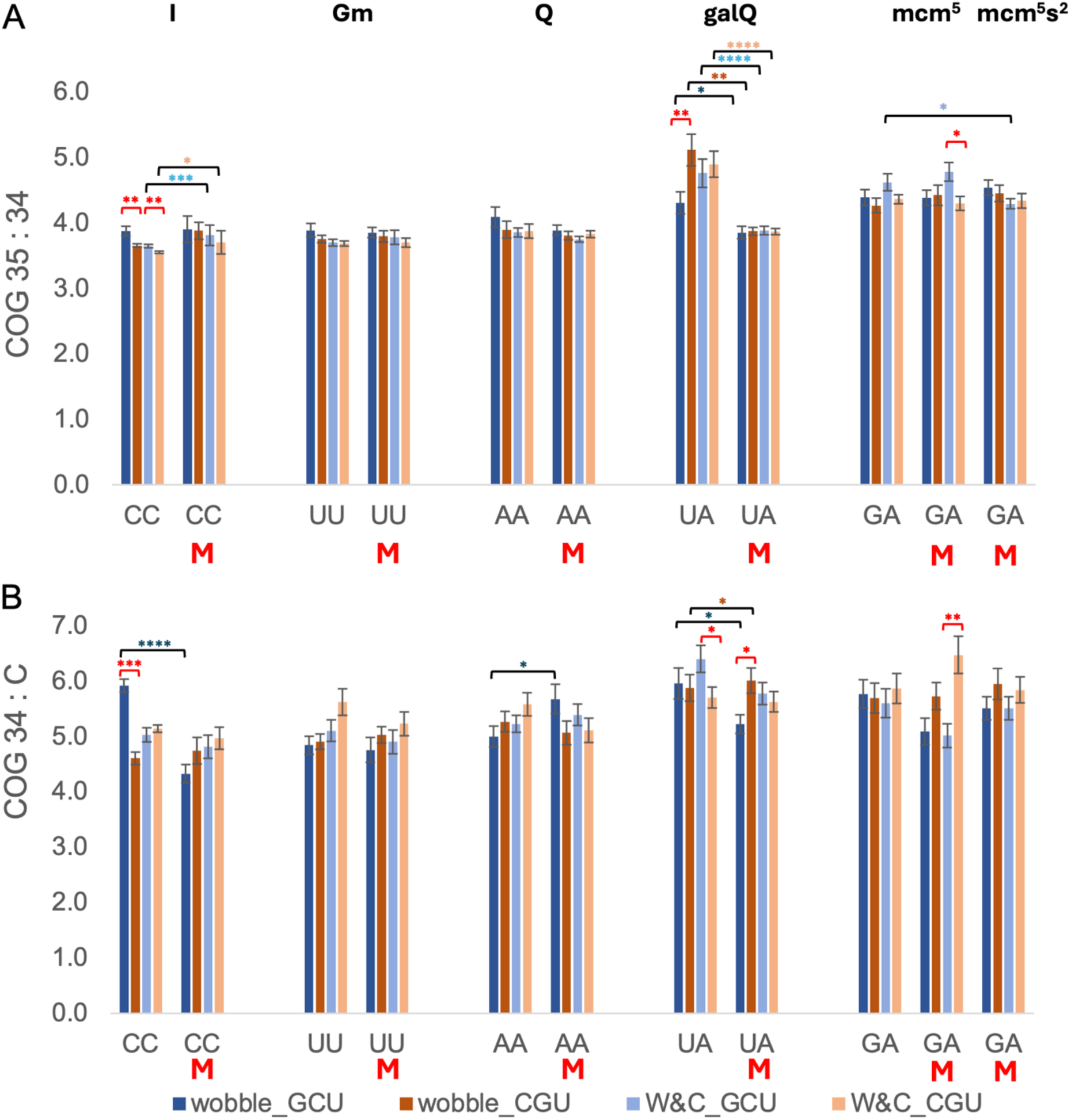
Modifications affect pi-pi stacking. Base stacking was assessed by measuring the distances between the centers of geometry (COG) of base rings^3^. (A) COG distance between nucleotide 35 and 34 of the tRNA anticodon. (B) COG distance between nucleotide 34 and the cytosine (C) of the CAR interaction surface. Student’s t-test p-values are denoted as: * p<0.05, ** p<0.01, *** p<0.001, **** p<0.0001, ns p≥0.05 (no * sign). Black brackets compare modified and unmodified nucleotide 34 structures; red brackets compare +1 GCU with +1 CGU structures,

In addition, we observed significant changes for I34_CC and mcm^5^s^2^U34_GA but specifically only for Watson-Crick H-bonding of nucleotide 34. Thus, tRNA anticodon stacking appeared to be influenced by the base-pair H-bonding of the tRNA with the mRNA, consistent with our previous observations^7^. The mcm^5^s^2^ effects were further restricted to the case where the CAR site +1 codon sequence is GCU, reflecting the influence of the CAR site on the tRNA anticodon.

We also assessed whether nucleotide 34 modifications influence the effects of the +1 codon on 34 : 35 stacking. Figure 4A (red brackets) highlights CAR site effects through student’s t-test comparisons of MD trajectories with +1 GCU or +1 CGU codons. These comparisons indicated significant effects of the +1 codon sequence on nucleotides 35 : 34 stacking, but interestingly only for several of the unmodified G34 structures (A site codons CCU, CCC and UAU; Fig. 4A) and modified mcm^5^U34 (A-site GAA). These results support the hypothesis that the modifications can tune the effects of the CAR site on the tRNA anticodon 35 : 34 stacking.

We further examined the relationship between the A-site and the adjacent CAR site by assessing COG distances between nucleotide 34 and C of CAR. When the tRNA nucleotide 34 base stacks with the C nucleotide of the CAR interface, it anchors CAR as an extension of the anticodon. Indeed, the 35 : 34 : C stacking has previously been highlighted as a potential structural communication channel between the A and CAR sites^7^. We found that the nucleotide 34 modifications altered the 34 : C stack and therefore the anchoring of CAR to the A site (Fig. 4B, black brackets). These effects were pronounced for the wobble +1 GCU background with the modifications I34_CC, Q34_AA and galQ34_UA. We also found that the nucleotide 34 modifications influence the effects of the +1 codon on 34 : C stacking (Fig 4B, red brackets). Significant effects were observed for unmodified (A site codons CCU and UAC) and modified galQ (A site UAU) and mcm^5^U34 (A site GAA), supporting the hypothesis that the modifications can tune the effects of the CAR site on anchoring through C : 34 stacking.

### Nucleotide 34 methylation tunes CAR braking

Our observations that nucleotide 34 modifications had effects on anchoring of CAR to the A-site anticodon led us to assess more generally whether the modifications affected behavior of the CAR surface, and more specifically the hypothesized braking behavior of CAR mediated by H-bonding to +1 GNN codons.

We assessed the H-bonding between CAR and the mRNA +1 codon (Fig. 5A). In all cases we always observed that the +1 GCU codon has stronger H-bonding with CAR than +1 CGU which reiterates the sequence dependency of the CAR functionality^2,3^. Here we refer to this difference in +1 GCU and +1 CGU behaviors as the CAR functionality range. To quantitatively assess this dynamic range, we normalized the H-bond values for each of the 30 +1 GCU MD simulation replicates by subtracting the H-bond average across all of the 30 +1 CGU replicates 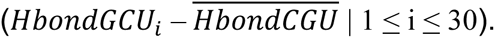 (Fig. 5A). The mean of these 30 normalized values (Fig. 5B: CAR site H-bonds (GCU-CGUavg)) provides a measure of the CAR H-bonding functionality range.

**Figure 5:**
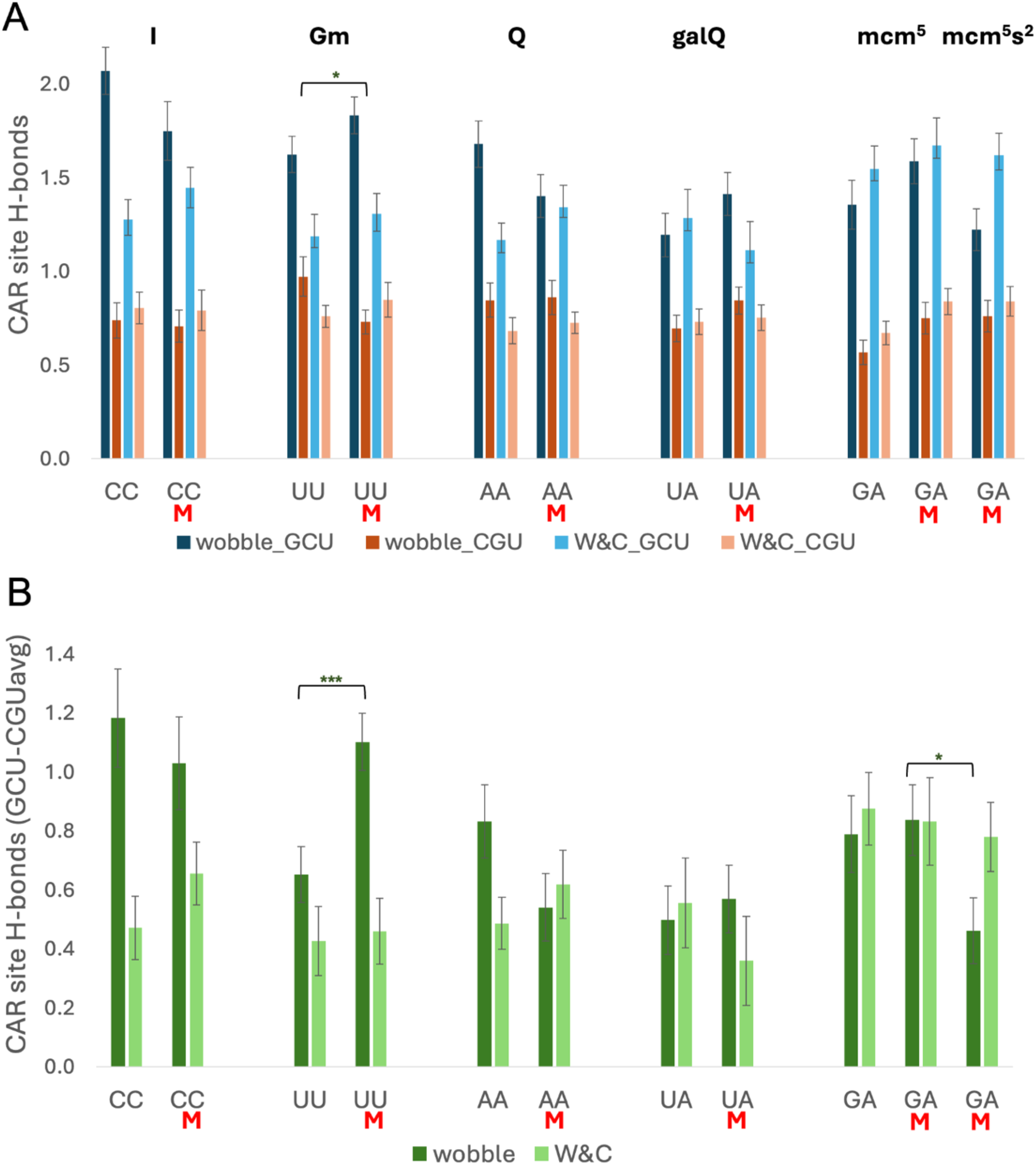
Modifications affect CAR site H-bonding. (A) Average H-bonding levels between the CAR residues and the mRNA +1 codon sequences GCU and CGU for different A-site and nucleotide 34 modification contexts. (B) CAR site H-bonds (GCU-CGUavg) shows the CAR functionality range (see text). Student’s t-test p-values are denoted as: * p<0.05, ** p<0.01, *** p<0.001, **** p<0.0001, ns p≥0.05 (no * sign).

Nucleotide 34 modifications can influence CAR by altering its functionality range (Fig. 5B). The most pronounced effect was observed for Gm34_UUU. When G34 is methylated to Gm34, the CAR functionality range significantly increases for the wobble A site UUU but not for the Watson-Crick A site UUC. Interestingly, Gm34 modification levels are increased under oxidative stress effectively storing information about the cellular environment ^33,34^. Since Gm34 methylation modulates the CAR functionality range, this suggests that oxidative stress may regulate translation in part through nucleotide 34 modification and its influence on the CAR site.

To investigate potential stress-mediated roles of CAR in translation of UUU codons, we performed dicodon analysis of published ribosome profiling data^10^ comparing ribosome densities when mapped to A-site UUU codons with or without +1 GNN. In line with our previous observations^3^, we observed higher ribosome densities for UUU with +1 GNN (UUUwGNN) dicodons, indicating slower translation, compared with UUU without +1 GNN (UUUwoGNN) (Fig 6A). Measurement of log_2_(UUUwGNN/UUUwoGNN) showed that this density difference was more pronounced for cells under oxidative stress (treated with 0.03% H_2_O_2_; Fig 6A, lower graph). This was observed for stressed cells in two large-scale profiling experiments (oxi1 and oxi2, Fig. 6A) which by bootstrap analysis had higher ratios than one (the larger, wo1) of the two independent profiling experiments with unstressed cells. This analysis parallels our analysis above suggesting that the more pronounced translation regulation in response to +1 GNN codons may in part be explained by stress-induced Gm34 methylation affecting CAR tuning of translation. In contrast to UUU, the UUC codon which base pairs with the same Phe 5’GmAA anticodon, but uses Watson-Crick base pairing at the wobble position, did not show an increase in the ribosome density ratio (UUCwGNN/UUCwoGNN) suggesting that wobble base pairing at the third codon position may contribute to the observed stress sensitivity (Figs. 5B, 6A).

**Figure 6:**
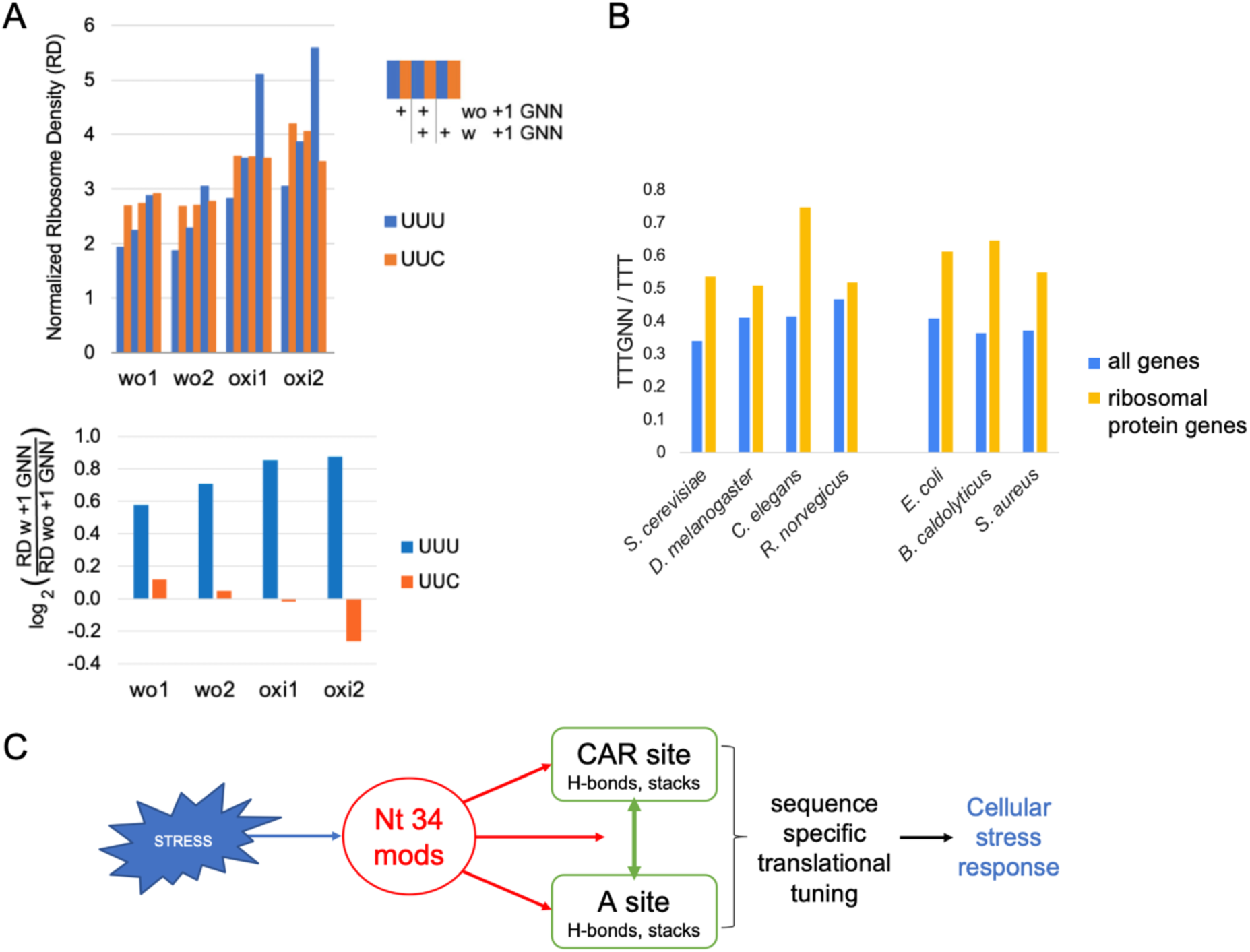
CAR mediated translational tuning and stress. (A) Dicodon analysis of published ribosome profiling data shows ribosomal densities (RD) at A-site UUU and UUC codons adjacent to +1 GNN or +1 (notG)NN codons. Two replicate experiments with unstressed yeast cells (wo1, wo2) were compared with cells exposed to oxidative stress (oxi1, oxi2) (upper graph). Assessing log_2_(RD w +1 GNN/RD wo +1 GNN) (lower graph) showed that UUU codons followed by +1 GNN exhibited higher ribosome densities, indicating slower translation, an effect that was enhanced in stressed cells. (B) Open reading frames (ORFs) in several species were analyzed to calculate the fractions of TTT codons that are followed by GNN codons for all genes (blue) and ribosomal protein genes (yellow). (C) Hypothetical model of sequence specific translational regulation under stress. Stress (blue arrow) dynamically regulates the levels of nucleotide 34 modifications which influence (red arrows) the functionality of A site, CAR site and their interplay (green arrows). The magnitudes of sequence specific CAR braking, as well as CAR mediated tuning of the A-site codon recognition interactions, alter under stress through the structural influences of nucleotide 34 modifications.

We explored further the hypothesis that UUUGNN dicodons might play a role in translation stress responses by examining the frequencies of these dicodons in ribosomal protein gene ORFs given that it is particularly important to downregulate expression of these highly-expressed genes under stress conditions to conserve energy^82–86^. Using sequence walkers to measure the fraction of UUU codons followed by adjacent GNN codons, we found in multiple species that UUUGNN dicodon frequencies are significantly elevated in ribosomal protein genes compared to all genes (of each species; Fig. 6B), consistent with the possibility that their evolutionary selection could aid in facilitating CAR-mediated depression of translation levels under stress. However, we note that GNN codon enrichment in ribosomal protein genes is observed for many yeast codons that utilize G34 tRNAs (not shown), including 3’ adjacent to UUC codons that use the same Phe 5’GmAA tRNA as UUU, suggesting that GNN enrichment in high-expression ORFs likely extends to mechanisms beyond the possible stress-response benefits.

In summary, our observations support a model in which the stress induced methylation of G34 signals CAR to expand its functionality range. This increases the difference in H-bonding of CAR to +1 GNN codons compared to +1 not GNN. Since this H-bonding is hypothesized to act as a brake that lowers the rate of translation, genes with higher frequencies of strongly binding +1 GNN codons are likely to be more susceptible to downregulation of translation under stress mediated in part by CAR. Thus, CAR is likely to act as a regulator of translation promoting differential gene expression in response to stress. One such cellular stress response mechanism appears to be selective downregulation of the ribosomal proteins to reduce ribosome biogenesis and global translation. As protein synthesis is a very energetically expensive cellular process, reducing the global translation levels could help the cells conserve energy under stressed conditions, thereby promoting survival. Since CAR is conserved across species, as is the selection for UUUGNN dicodons in ribosomal protein genes, this hypothesized regulatory mechanism has likely been conserved through evolution.

In our analysis of ribosome profiling data, we focused on CAR functionality and the differences in translation rates for dicodons with and without GNN. However, assessing translation rates without reference to GNN codons revealed that under oxidative stress, UUU and UUC codons have higher ribosome densities and therefore slower translation than other codons (Fig. 6A, upper graph). This result lies in contrast to previous suggestions that translation of these Phe codons is enhanced under oxidative stress. Indeed, a study on mouse brain cells showed that cells with knock out of Gm34 methyltransferase Ftsj1, resulting in Gm34 loss, leads to translation slowdown at UUU and UUC codons, particularly in neurons, linking Gm34 to intellectual disability^41^. However, these results could be partially explained by tRNA abundance effects. Several other studies involving humans, flies and mice have established the clinical importance of Gm34 in human pathology by correlating it with neuronal structure, cognitive function, X-linked intellectual disability and also cancer metabolism^31,37–44^.

## Conclusions

Our findings suggest a model that stages CAR as a stress-sensitive regulator of translation (Fig. 6C). Cells exist under dynamically varying environmental conditions. They constantly sense and respond to the environmental cues like oxidative stress. Here we propose a layer of the cellular stress response based on analysis of the effects of nucleotide 34 modifications that are regulated by stress. These stress-sensitive chemical modifications alter H-bonding and stacking interactions of the A and CAR sites which in turn are associated with tuning of translational fidelity and speed. Moreover, both the A and CAR sites engage in a mutual communication network where each site influences the structure of the other and this communication is sensitive to modifications of tRNA nucleotide 34 leading to altered responses to +1 codons at the CAR site.

The prominent role of +1 GNN codons adjacent to stress-sensitive A-site codons such as UUU appears to have led to selection of GNN dicodon enrichment in gene sets whose expression is reprogrammed under stress environmental conditions. The dynamic nature of the reprogramming mediated by controlling tRNA modifications likely provides one of the mechanisms for phenotypic plasticity in response to changing environments.

## ASSOCIATED CONTENT

Supporting information:

MD restart and topology files are available at https://doi.org/10.25438/wes02.32411568;

Analysis code can be found at https://github.com/mitsuraval/nt34_modifications and Sun et al.^3^;

## AUTHOR INFORMATION

## Author contributions

The manuscript was written through contributions of all authors. All authors have given approval to the final version of the manuscript.

## Funding Sources

This work was supported by the National Institutes of Health [GM120719 to M.P.W., GM128102 to K.T.] and the National Science Foundation [CHE-2320718 to the MERCURY consortium]. Funding for open access charge: National Institutes of Health.

## ACKNOWLEDGEMENT

We thank Joseph Coolon, Scott Holmes, Phil Arevalo and Amy MacQueen for discussions, and Henk Meij for technical assistance with high-performance computing. We acknowledge the use of Generative AI (ChatGPT version 4o) for text copy editing and to enhance code where appropriate.

## Notes

https://doi.org/10.25438/wes02.32411568

### Competing Interest Statement

The authors have declared no competing interest.

