## Supplemental figures and tables for "Anticodon nucleotide modifications affect translational tuning by the ribosomal CAR surface"

#### Supplementary Figures:

| | Trajectory length | Unmodified structure | Modified structure |
|---|---|---|---|
| AAUGCU | 60ns | 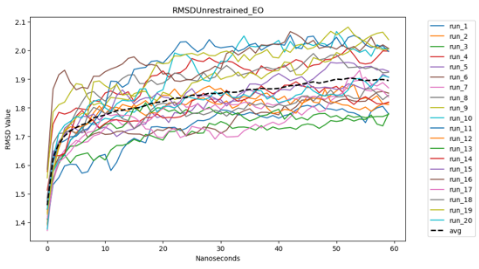  | 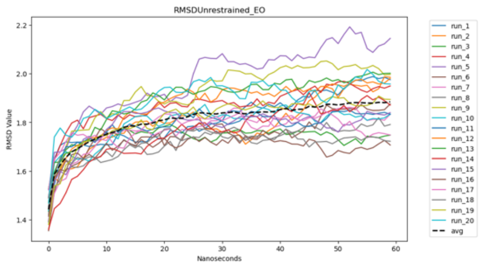  |
| | 100ns | 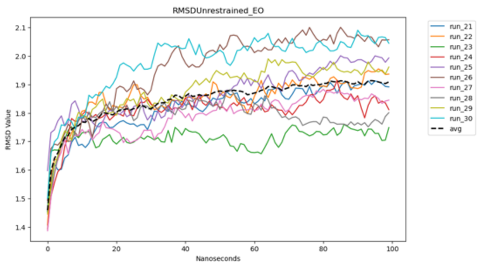  | 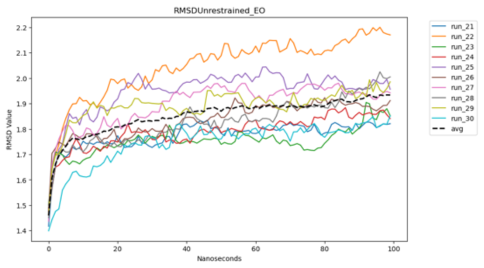  |

<table>
<tr><td rowspan="2">AAUCGU</td><td>60ns</td><td>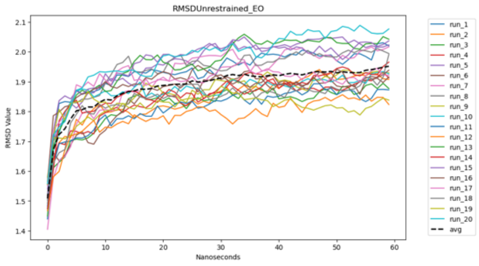
</td><td>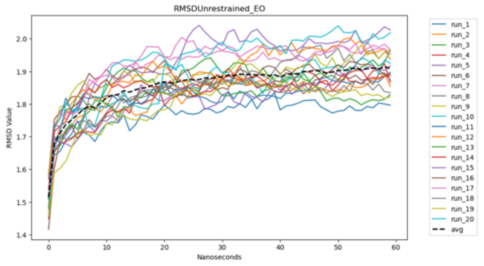
</td></tr>
<tr><td>100ns</td><td>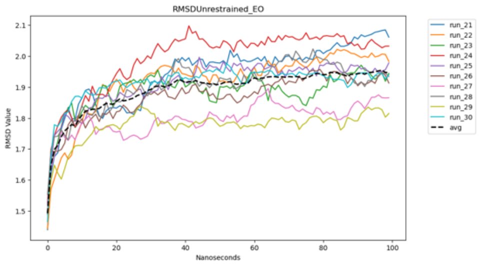
</td><td>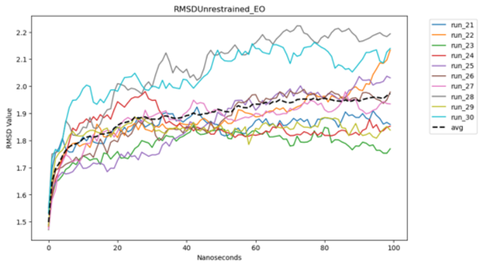
</td></tr>
<tr><td rowspan="2">AACGCU</td><td>60ns</td><td>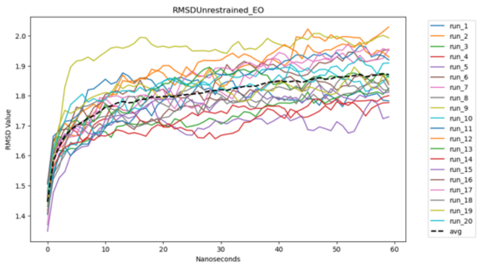
</td><td>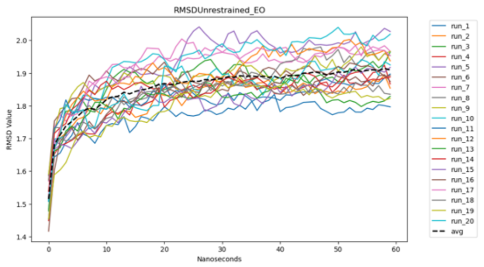
</td></tr>
<tr><td>100ns</td><td>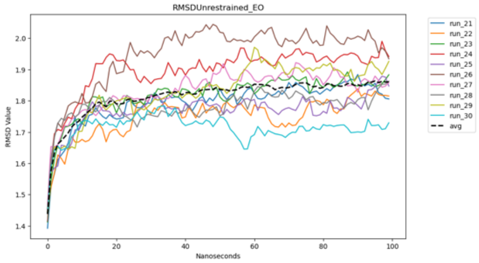
</td><td>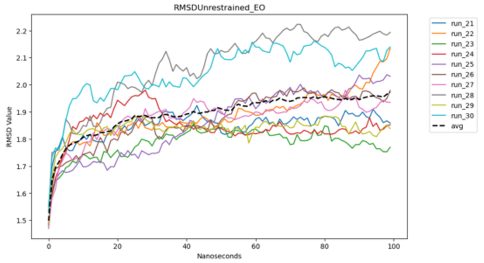
</td></tr>
<tr><td>AACCGU</td><td>60ns</td><td>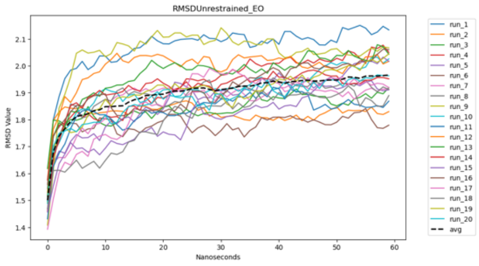
</td><td>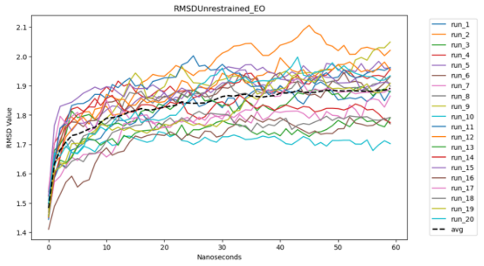
</td></tr>
</table>

| | | | |
|---|---|---|---|
| | 100ns | 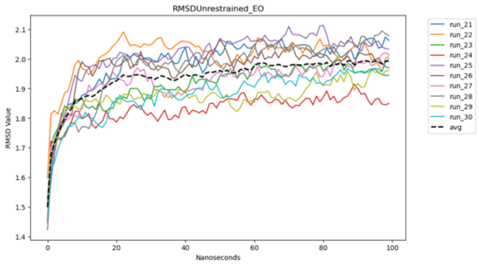 | 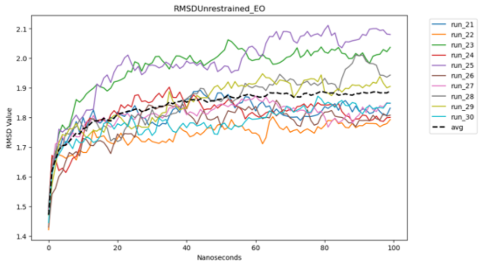 |





Figure S1: RMSD values with reference to equilibration output structures for backbone atoms of unrestrained residues across 0-60ns or 0-100ns trajectories shows settled dynamics after the first 20ns of the trajectories.

**Supplementary Table:**

**Table S1**

| # | modification status | A-site N1 N2 | A-site N3 | +1 codon |
|---|---|---|---|---|
| 1 | G34 | CC | U (wobble) | GCU |
| 2 | G34 | CC | U (wobble) | CGU |
| 3 | G34 | CC | C (W&C) | GCU |
| 4 | G34 | CC | C (W&C) | CGU |
| 5 | I34 | CC | U (wobble) | GCU |
| 6 | I34 | CC | U (wobble) | CGU |
| 7 | I34 | CC | C (W&C) | GCU |
| 8 | I34 | CC | C (W&C) | CGU |
| 9 | G34 | UU | U (wobble) | GCU |
| 10 | G34 | UU | U (wobble) | CGU |
| 11 | G34 | UU | C (W&C) | GCU |

| | | | | |
|---|---|---|---|---|
| 12 | G34 | UU | C (W&C) | CGU |
| 13 | Gm34 | UU | U (wobble) | GCU |
| 14 | Gm34 | UU | U (wobble) | CGU |
| 15 | Gm34 | UU | C (W&C) | GCU |
| 16 | Gm34 | UU | C (W&C) | CGU |
| 17 | G34 | AA | U (wobble) | GCU |
| 18 | G34 | AA | U (wobble) | CGU |
| 19 | G34 | AA | C (W&C) | GCU |
| 20 | G34 | AA | C (W&C) | CGU |
| 21 | Q34 | AA | U (wobble) | GCU |
| 22 | Q34 | AA | U (wobble) | CGU |
| 23 | Q34 | AA | C (W&C) | GCU |
| 24 | Q34 | AA | C (W&C) | CGU |
| 25 | G34 | UA | U (wobble) | GCU |

| | | | | |
|---|---|---|---|---|
| 26 | G34 | UA | U (wobble) | CGU |
| 27 | G34 | UA | C (W&C) | GCU |
| 28 | G34 | UA | C (W&C) | CGU |
| 29 | galQ34 | UA | U (wobble) | GCU |
| 30 | galQ34 | UA | U (wobble) | CGU |
| 31 | galQ34 | UA | C (W&C) | GCU |
| 32 | galQ34 | UA | C (W&C) | CGU |
| 33 | G34 | GA | A (wobble) | GCU |
| 34 | G34 | GA | A (wobble) | CGU |
| 35 | G34 | GA | G (W&C) | GCU |
| 36 | G34 | GA | G (W&C) | CGU |
| 37 | $mcm^5U34$ | GA | A (wobble) | GCU |
| 38 | $mcm^5U34$ | GA | A (wobble) | CGU |
| 39 | $mcm^5U34$ | GA | G (W&C) | GCU |

| | | | | |
|---|---|---|---|---|
| 40 | $mcm^5U34$ | GA | G (W&C) | CGU |
| 41 | $mcm^5s^2U34$ | GA | A (wobble) | GCU |
| 42 | $mcm^5s^2U34$ | GA | A (wobble) | CGU |
| 43 | $mcm^5s^2U34$ | GA | G (W&C) | GCU |
| 44 | $mcm^5s^2U34$ | GA | G (W&C) | CGU |

### Forty-four structures were assessed by MD in this study. All 44 structures have unique combinations of variables. The variables are (1) modification status and corresponding A-site nucleotides 1 and 2 (N1 N2), (2) Wobble or Watson-Crick base pairing determined by A-site nucleotide 3 (N3) identity, and (3) the +1 codon sequence GCU or CGU.
